# The Effect of New Nurses’ Clinical Competence on Career Adaptation

**DOI:** 10.1101/2019.12.20.884270

**Authors:** Kyu Ho Lee, Mi Joon Lee

## Abstract

**Background:** New nurses need a certain period of time to adapt to the organization due to a lack of clinical competence, and also immaturity in social skill. The purpose of this study was to evaluate the effects of new nurses’ clinical competence on career adaptation in order to use the results as basic information for developing education programs to improve their adaptability in clinical practice.

**Method:** This study employed a descriptive survey to investigate the clinical competence and the career adaptability of new nurses working in a general hospital. The study subjects were 61 new nurses with less than one year of work experience and data was collected from June, 2018 to July, 2019. Data was analyzed using frequency analysis, descriptive statistics, t-test, ANOVA, Pearson’s correlation coefficients, and multiple regression analysis.

**Results:** On average, the study subjects have worked for 11.33±1.51 months. In this study, the average clinical competence of new nurses was 2.21±0.61, and career adaptability was 3.00±0.39. The clinical competence of new nurses was positively correlated with career adaptability (r=.453, p<.001). Among the sub-categories of clinical competence, developing professional and legal implementation were found to affect career adaptability (t=2.24, p=.030).

**Conclusion:** The clinical competence of the new nurses was shown to positively affect their career adaptability, and it was confirmed that developing professional and legal implementation in the subcategories of clinical competence affected the career adaptability of the new nurses. Thus, it is necessary to establish a program that provides support for new nurses to enable them to build professional identities that they can be proud of.

## Introduction

With the recent advances in medical technology and the improvement of people’s living and educational standards, interest in quality medical services, as well as demand for high quality nursing care, have increased. Accordingly, nurses must acquire sufficient medical knowledge and competent skills [1]. However, new nurses are facing many challenges in the fast-paced hospital environment while also experiencing high stress and maladjustment [2].

New nurses are those who have graduated from a nursing college in Korea and obtained a nurse’s license with working experience of less than 12 months after being employed in a medical institution [3]. New nurses face a lot of stress in their clinical careers after their first year of graduation as they have to deal with the gap between high expectations and reality in the following aspects: their practical skills; demand for knowledge and skills required for nursing work; three-shift working environment; increased responsibility for nursing behavior; subtle interpersonal relationships with colleagues; and formation of interpersonal relationships with patients and caregivers [4, 5]. In addition, new nurses need a certain period of time to adapt to the organization due to lack of clinical competence, difficulty in making their own decisions, and immaturity in career adaptability [6].

Clinical competence means the ability to perform the task of obtaining the expected results in the practical working environment, and new nurses are required to secure necessary clinical competence for them to grow into professional nurses and adapt to clinical practice [7]. However, it has been pointed out that the clinical competence of graduated nurses does not meet the demands of the nursing field in the hospital setting, and the maladaptation of new nurses to the nursing field due to lack of clinical competence has a negative effect on job satisfaction and professional identity among clinical nurses, resulting in turnover [3]. The turnover of new nurses leads to a loss of management expense for medical personnel [8] and increases the overtime work and stress of in-service nurses, making them unable to meet the need of patients for quality care from skilled nursing manpower. Thus, multidimensional efforts have been made to analyze the factors related to the clinical competence of nurses and to develop and apply educational programs for improving the clinical competence of nurses in the clinical setting.

Career adaptability is the process of accepting the established policy of an organization to which an individual belongs, internalizing the organization’s norms and values, and forming a professional identity through the work-related knowledge, skills acquisition, and supportive interrelationships that the organization needs to transform into a productive member of the organization [9]. However, new nurses are overwhelmed by psychological pressures, tensions, and fears associated with the characteristics of nursing work, and they easily make mistakes or errors due to their inexperience. As a result, they lose their self-confidence and self-esteem and experience negative emotions such as a victim mentality or a sense of guilt as they are accused and criticized by their fellow nurses. Since the continuing psychological stress from work leads to serious interpersonal conflict and difficulties in career adaptability [6], efforts are needed to promote the career adaptability of new nurses.

In Korea, previous studies on the clinical competence of new nurses include predictors of clinical competence in new graduate nurses [7], the relationship between clinical competence and field adaptation [10], the effect of a practical work-oriented education program on the ability of newly recruited nurses in execution of clinical competency [11], the relationship between critical thinking disposition and clinical competence [12], etc. Related studies on career adaptability are as follows: factors related to organizational socialization of new nurses [6], influence of workplace bullying and resilience on organizational socialization [13], resilience and organizational socialization [14], professional self-concept and organizational socialization [15], and the effect of a new nurse-preceptor exchange relationship on organizational socialization [16]. Moreover, clinical competence and degree of organizational socialization were studied in relation to the communication style of preceptors [17] and teaching style of preceptors [18]. On the other hand, there were a few studies that researched clinical competence and career adaptability independently.

The purpose of this study was to explore ways to improve clinical competence and career adaptability of new nurses by identifying the degree of clinical competence and career adaptability perceived by new nurses and determining the relationship among them. The specific objectives were as follows.

## Materials and Methods

### Subjects

This descriptive study was designed to identify clinical competence and career adaptability as perceived by new nurses working in a general hospital and it also examined the correlation between variables.

The subjects of this study were new nurses with less than a year of work experience in general wards, intensive care units, and emergency rooms and those who provided direct care to patients in their assigned service department for less than a year.

They agreed in writing to participate in the research. Excluded in this study were the nurses who did not agree to participate or those who did not provide direct care to patients such as those working in operating rooms, anesthesia departments, hemodialysis units, and endoscopic clinics.

### Instruments

#### Clinical competence

For measurement of clinical competence, the instrument that was developed by Son et al. [20] to assess the basic clinical nursing competence of general nurses, regardless of particular medical fields, was implemented based on the educational objectives of core nursing competence-centered nursing practices that was proposed by Kim [19]. This tool, which encompasses the seven core competencies, consists of a total of 64 questions on clinical competence including five questions on data collection, 24 questions on basic nursing care, six questions on communication, six questions on critical thinking, nine questions on teaching and leadership, 11 questions on nursing management, and five questions on developing professional and legal implementation. Each question has a four-point scale, ranging from 1 point for ‘Cannot do at all’ to 4 points for ‘Can do very well.’ Higher scores indicate higher clinical competence. In the study conducted by Son et al. [20], the Cronbach’s α was 0.82, and that of this study was 0.99.

#### Career adaptability

Career adaptability refers to the process of learning the values, expected performance and behavior, and social knowledge that are needed to perform one’s role in an organization as a member [21]. Career adaptability was measured using 39 questions on career adaptability of new nurses developed by Sohn et al. [2]. This instrument consists of eight questions on personal characteristics, eight questions on organizational characteristics, three questions on professional identity, five questions on job performance, five questions on job satisfaction, five questions on organizational commitment, and five questions on burnout. Each question has a five-point scale, ranging from 1 point for ‘Never’ to 5 points for ‘Always’. Higher scores indicate higher career adaptability. In the study conducted by Sohn et al. [2], the Cronbach’s α was 0.97, and that of this study was 0.85.

### Data collection

This study was conducted with 61 new nurses who were hired in 2017 and the survey was conducted from June 2018 to July 2019. After the researcher explained the purpose, necessity, and procedure of the study directly to the study subjects, the questionnaire was distributed to those who voluntarily agreed to participate in the study through written consent. The questionnaire was submitted directly to a collection box with an installed lock at a separate location.

### Statistical analysis

The data was analyzed using IBM Statistics 24 program and the two-tail test was conducted at 95% level of confidence to determine statistical significance.

The general characteristics, clinical competence, and degree of career adaptability among the subjects were analyzed through frequency, percentage, mean, and standard deviation. Clinical competence and career adaptability depending on the subjects’ general characteristics were analyzed by t-test and ANOVA, followed by Scheffe’s test as a post-hoc analysis. The correlation between the subjects’ clinical competence and career adaptability was analyzed by Pearson’s correlation coefficients. In addition, the effect of clinical competence on career adaptability was analyzed using multiple regression analysis. The statistical significance of the analysis result was determined by the two-tail test conducted at 95% level of confidence.

### Ethics consideration

This study was conducted after obtaining approval from the Institutional Review Board of the K General Hospital (2018-03-002-003), and a questionnaire collection box was installed at a separate place to maintain anonymity and autonomy.

## Results

The new nurses, who were the subjects of this study, were mostly women (85.2%) and the majority had obtained a bachelor’s degree (95.1%) (Table 1). On average, the duration of their work experience was 11.33±1.51 months. The departments where they worked were internal medicine wards (42.6%), surgery wards (34.4%), and special parts (23.0%). More than half of the subjects (65.6%) had been placed in their desired departments, while 57.4 % were trained in a medical department similar to their currently assigned department.

**Table 1.** General characteristics of new nurses (*N*=61)

| Variable | Category | Value |
| --- | --- | --- |
| Age (years) | | $24.48 \pm 1.70$ |
| Career Duration (months) | | $11.33 \pm 1.51$ |
| Gender | Male | 9 (14.8%) |
|  | Female | 52 (85.2%) |
| Education | College | 3 (4.9%) |
|  | University | 58 (95.1%) |
| Working department | Internal medicine ward | 26 (42.6%) |
|  | Surgery Ward | 21 (34.4%) |
|  | Special Part | 14 (23.0%) |
| Desired department | Yes | 40 (65.6%) |
|  | No | 21 (34.4%) |
| Experience of working department | Yes | 35 (57.4%) |
|  | No | 26 (42.6%) |

Values are presented as n (%) or mean±standard deviation

The degree of clinical competence perceived by the new nurses was 2.21±0.61 points on average. Teaching and leadership (2.28±0.59 points) and nursing management (2.28±0.54 points) showed the highest scores among the seven subcategories of clinical competence, followed by developing professional and legal implementation (2.25±0.62 points), critical thinking (2.21±0.71 points), data collection (2.18±0.69 points), and basic nursing care (2.17±0.70 points). Communication was found to be the lowest with 2.14±0.77 points (Table 2).

**Table 2.** Perception of clinical competence (*N*=61)

| Category | Min | Max | Mean±SD |
| --- | --- | --- | --- |
| Data collection | 1.00 | 4.00 | $2.18 \pm 0.69$ |
| Basic nursing care | 1.00 | 4.00 | $2.17 \pm 0.70$ |
| Communication | 1.00 | 4.00 | $2.14 \pm 0.77$ |
| Critical thinking | 1.00 | 4.00 | $2.21 \pm 0.71$ |
| Teaching & leadership | 1.22 | 3.89 | $2.28 \pm 0.59$ |
| Nursing management | 1.09 | 3.91 | $2.28 \pm 0.54$ |
| Developing professional & legal implementation | 1.00 | 4.00 | $2.25 \pm 0.62$ |
| Clinical Competence average | 1.14 | 3.75 | $2.21 \pm 0.61$ |

SD=standard deviation

The degree of career adaptability perceived by the new nurses was 3.00±0.39 points in average. Burnout (3.40±0.82 points) showed the highest score among the seven subcategories of career adaptability, followed by organizational commitment (3.17±0.62 points), organizational characteristics (2.99±0.57 points), job satisfaction (2.94±0.48 points), personal characteristics (2.68±0.84 points), and professional identity (2.64±0.78 points). Job performance was found to be the lowest with 2.11±0.90 points (Table 3).

**Table 3.**
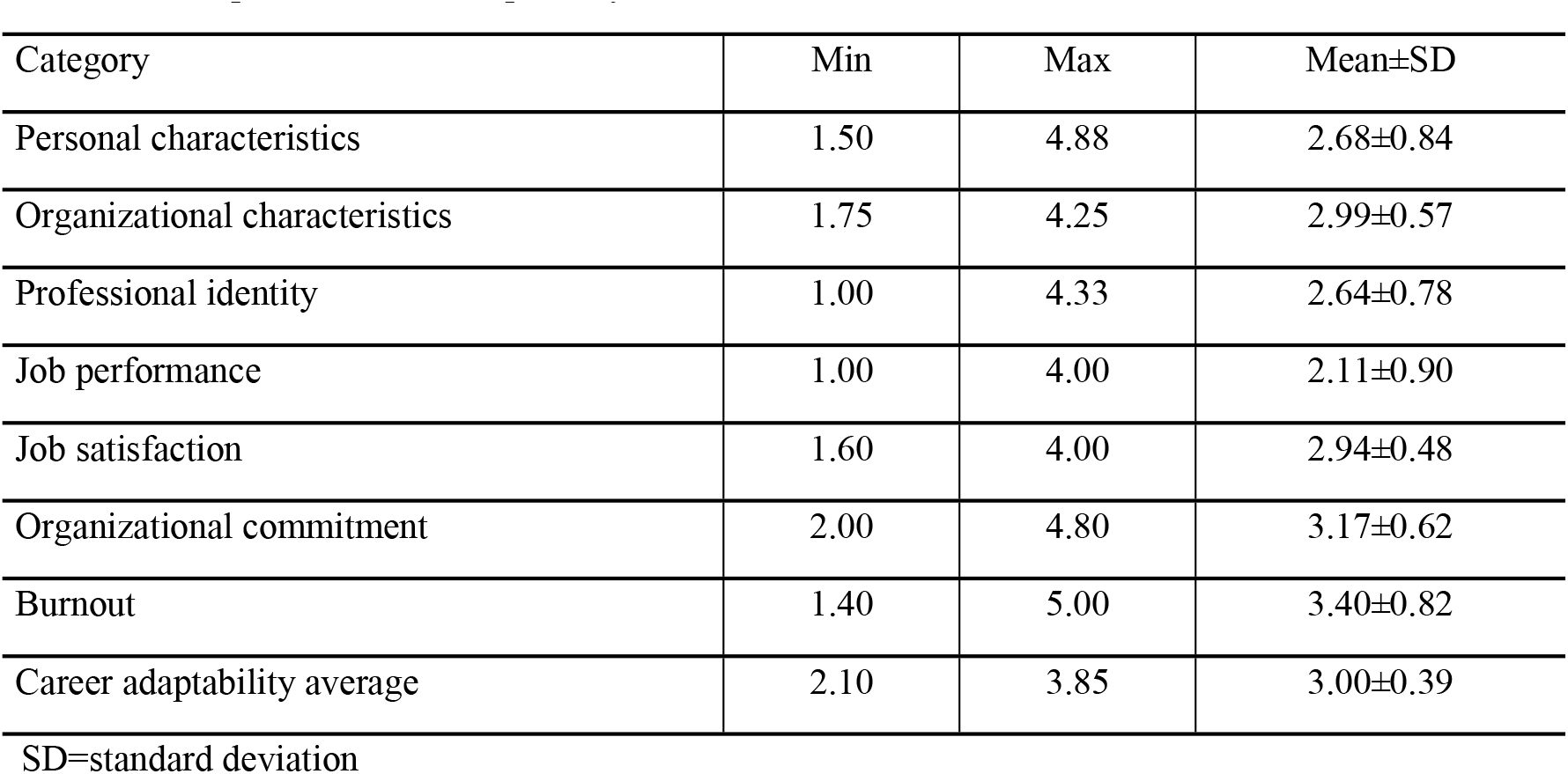
Perception of career adaptability (*N*=61)

| Category | Min | Max | Mean±SD |
| --- | --- | --- | --- |
| Personal characteristics | 1.50 | 4.88 | 2.68±0.84 |
| Organizational characteristics | 1.75 | 4.25 | 2.99±0.57 |
| Professional identity | 1.00 | 4.33 | 2.64±0.78 |
| Job performance | 1.00 | 4.00 | 2.11±0.90 |
| Job satisfaction | 1.60 | 4.00 | 2.94±0.48 |
| Organizational commitment | 2.00 | 4.80 | 3.17±0.62 |
| Burnout | 1.40 | 5.00 | 3.40±0.82 |
| Career adaptability average | 2.10 | 3.85 | 3.00±0.39 |

SD=standard deviation

Regarding clinical competence depending on general characteristics of the new nurses, there were no statistical differences in sex (t=0.82, p=.435), education (t=−0.94, p=.351), working department (F=0.34, p=.712), desired part (t=0.05, p=.958), and experience in working department (t=0.39, p=.699). In terms of career adaptability, it was found that there were no statistical differences in terms of sex (t=0.56, p=.580), education (t=−0.63, p=.528), working department (F=1.80, p=.175), desired part (t=−0.16, p=.877), and experience in working department (t=−0.19, p=.852) (Table 4).

**Table 4.** General characteristics of clinical competence and career adaptability (*N*=61)

| Variables | Categories | Clinical competence |  | Career adaptability |  |
| --- | --- | --- | --- | --- | --- |
|  |  | Mean±SD | t or F(P) Scheffe | Mean±SD | t or F(P) Scheffe |
| Sex | Male | 2.41±0.84 | 0.82 (.435) | 3.07±0.53 | 0.56 (.580) |
|  | Female | 2.18±0.56 |  | 2.99±0.37 |  |
| Education | College | 1.89±0.20 | -0.94 (.351) | 2.86±0.47 | -0.63 (.528) |
|  | University | 2.23±0.62 |  | 3.01±0.39 |  |
| Working department | medical ward | 2.19±0.65 | 0.34 (.712) | 2.94±0.41 | 1.80 (.175) |
|  | Surgical Ward | 2.16±0.53 |  | 3.13±0.38 |  |
|  | Special Part | 2.33±0.66 |  | 2.93±0.34 |  |
| Desired part | Yes | 2.22±0.57 | 0.05 (.958) | 3.00±0.39 | -0.16 (.877) |
|  | No | 2.21±0.68 |  | 3.01±0.40 |  |
| Experience in working department | Yes | 2.24±0.60 | 0.39 (.699) | 3.00±0.40 | -0.19 (.852) |
|  | No | 2.18±0.63 |  | 3.01±0.38 |  |

SD=standard deviation

There was a positive correlation between new nurses’ clinical competence and career adaptability (r=.453, p<.001), implying that the higher the clinical competence, the higher the career adaptability (Table 5).

**Table 5.**
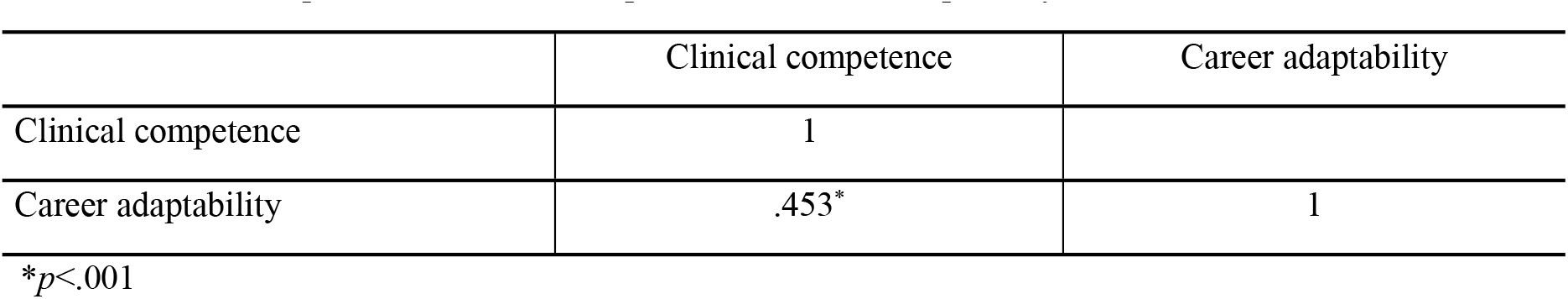
Relationship between clinical competence and career adaptability (*N*=61)

|  | Clinical competence | Career adaptability |
| --- | --- | --- |
| Clinical competence | 1 |  |
| Career adaptability | .453* | 1 |

\* $p<.001$

As for the factors that influence career adaptability in the subcategories of clinical performance as shown in Table 6, a 1-point increase in developing professional and legal implementation resulted in a 0.32-point increase in career adaptability, and as such was shown to be statistically significant (*p*=.030). This was shown to have 33.3% explanation power and was statistically significant (*p*=.002).

**Table 6.**
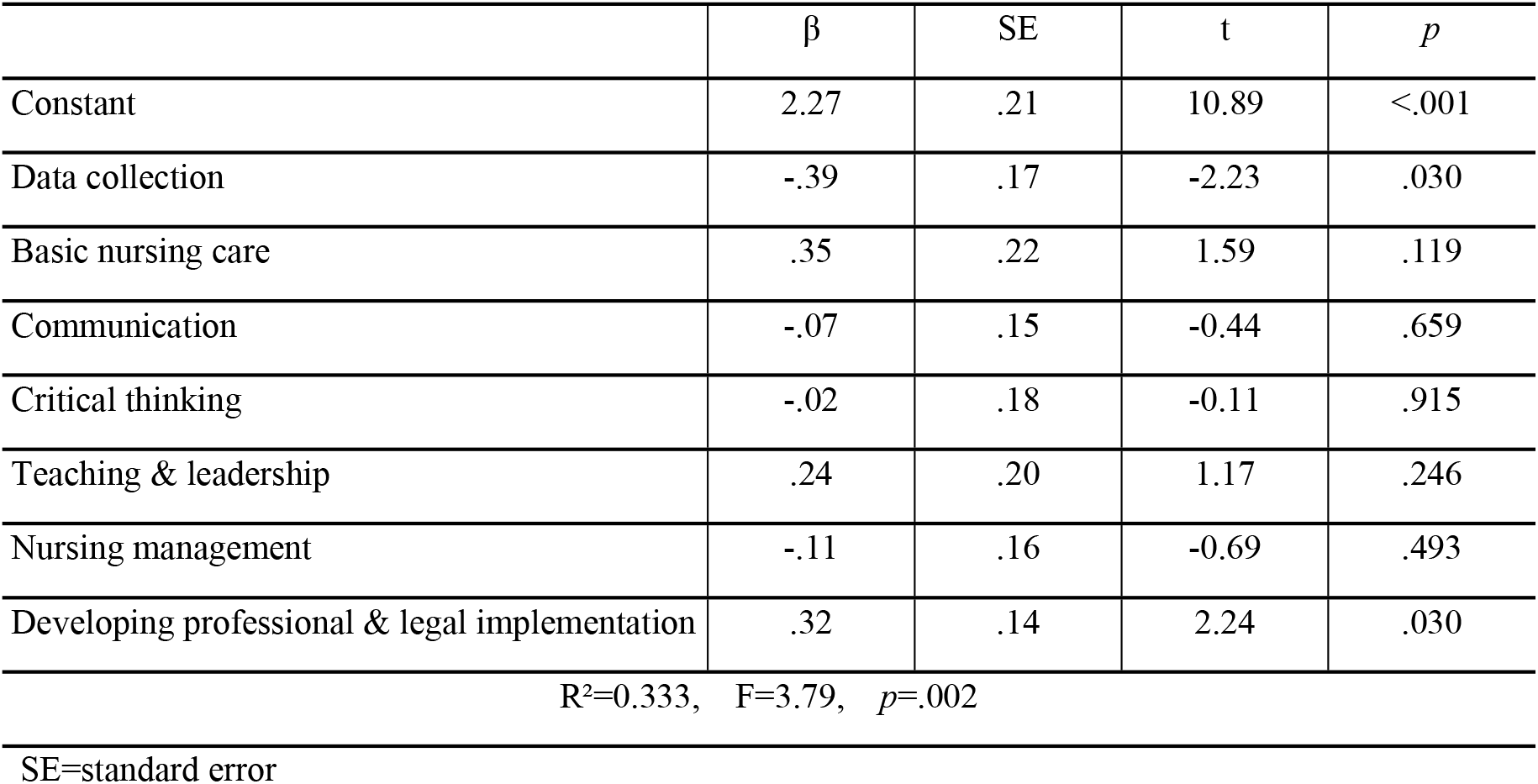
Effect of Clinical Competence on Career adaptability (*N*=61)

## Discussion

The purpose of this study was to examine the clinical competence and career adaptability as perceived by new nurses in general hospitals in order to use the results as basic data for practical application and program development to improve clinical adaptation of new nurses.

The average clinical competence perceived by the new nurses in this study was 2.21 points, which was lower than 2.82 points and 2.83 points, respectively obtained from the studies of Park and Park [17] and Shin et al. [7] using the same instrument. In previous studies, the average duration of work experience of new nurses was 7.2 months in the study of Park and Park [17] and 4.66 months in the study of Shin et al. [7], which were 11.33 months less than the average career duration obtained in this study. Given the differences in required nursing competence depending on career level, beginners with low clinical experience are expected to nurse patients with guidance or help in stable situations, while those with career duration of more than a year are expected to provide independent nursing services [22]. In addition, new nurses may also have difficulty in providing independent care in the higher-level nursing areas, and they may need an integrated ability to solve the patients’ changing needs based on their nursing knowledge, in addition to routine nursing activities or acquiring basic nursing skills in the course of their growth as a professional [7]. Therefore, it is believed that proper education and support should be provided to new nurses as their role as a practitioner increases.

In this study, teaching and leadership and nursing management were shown to have highest scores with 2.28 points among the subcategories of clinical competence, whereas communication had the lowest score with 2.14 points. This was in contrast to the results reported in the study of Park and Park, which showed the highest score in communication and lowest score in teaching and leadership, and the study of Shin et al. [7], which showed high scores in basic nursing care and communication and low scores in teaching and leadership and nursing management. According to the study conducted by Ji and Kim [23], functional nursing practice was the most important for new nurses with less than seven months of career duration, while those with more than a year of work experience were expected to serve in a professional role more often than those in basic nursing, such as taking on the role of a preceptor and developing educational materials [24]. These results may be attributed to the expanded communication opportunities among the nurses with about a year of work experience in and out of their professions when performing simple nursing tasks as their roles and experience with other departments were extended [25]. Therefore, specific educational programs to reduce conflicts in their roles, which may be caused by the expanded role of new nurses, and practical and case-specific concrete and diverse communication methods should be prepared.

In this study, the average career adaptability perceived by the new nurses was 3.00 points, and it was similar to the results obtained in previous studies: 3.03 points in the study of Kim and Yoo [26] and 3.20 points in the study of Kwak and Kwon [15]. In the subcategories of career adaptability, burnout and organizational commitment were higher with more than 3 points whereas job performance was the lowest with 2.11 points. Career adaptability means field adaptation and burnout in the subcategories includes reversed items; therefore, higher scores indicate low burnout perceived by the subjects. In this context, higher level of burnout and organizational commitment are considered to be higher career adaptability. In this study, burnout and organizational commitment in the subcategories of career adaptability resulted in higher scores and such results were similar to the outcome of the study conducted by Han et al. [27] that reported organizational commitment and burnout as the most important factors affecting the turnover of new nurses. In addition, job performance had the lowest scores in the subcategories of career adaptability, and it was similar to the results of the study conducted by Kwak and Kwon [15]. These were in line with the results reported in the study of Ji and Kim [23] that the most important factor affecting the turnover of new nurses with less than seven months of work experience was functional nursing competence; however, for new nurses with more than seven months of career duration, the most important factor was organizational commitment. Therefore, in order to enhance the career adaptability of new nurses, it is necessary to provide education that improves functional nursing competence (such as nursing techniques) at the early stages of the nurses’ career. Moreover, it is essential to institute measures that reduce burnout and increase organizational commitment among the nurses who provide independent nursing.

In this study, clinical competence and career adaptability were positively correlated (r=.453, p<.001). This was consistent with the result of the study conducted by Park and Park [17] that reported a statistically significant positive correlation between clinical competence and career adaptability (r=.24, p=.001). In particular, developing professional and legal implementation was shown to have effect among the subcategories of clinical competence (t=2.24, p=.030). According to the study of Kwak and Kwon [15], higher professional identity tended to show higher career adaptability and in the study of Son et al. [4], new nurses’ turnover intention was affected by professional nursing values and pride as a nurse. Such results were said to be consistent with the findings of this study. Therefore, it is necessary to come up with a program in which new nurses can form their professional identity and take pride as a nurse with a positive attitude while performing nursing tasks. In this study, however, the results of data collection factor among the subcategories of clinical competence (t=−2.23, p=.030) was unexpected and it may be necessary to verify the results through repeated studies and examination.

## Conclusions and recommendations

The purpose of this study was to examine the clinical competence and career adaptability as perceived by new nurses in general hospitals in order to use the results as basic data for practical application and program development to improve the clinical adaptation of new nurses. The study found that clinical competence and career adaptability, according to the general characteristics of the new nurses, did not show a statistically significant difference. The clinical competence of the new nurses was shown to positively affect their career adaptability, and it was confirmed that developing professional and legal implementation in the subcategories of clinical competence affected the career adaptability of the new nurses. Therefore, it is necessary to develop a program that helps form the professional identity and pride of new nurses.

The significance of this study may be found in the fact that it provided the basis for developing measures for career adaptability and field adaptation of new nurses by identifying specific factors affecting their career adaptability. Interpretation of the study results was limited as the study was conducted in a general hospital. Although complete enumeration was implemented to the new nurses, the study is considered to be limited due to the small number and convenient sampling of the subjects by the researcher. Therefore, it is necessary to conduct repeated surveys by increasing the number of subjects. In addition, it is recommended to: 1) develop and apply a program that helps in the career adaptability and field adaptation of new nurses based on improved clinical competence; and 2) conduct a study that verifies the effects of the program.

## Acknowledgements

The authors acknowledge the efforts of the nursing education group at Kangbuk Samsung Hospital, Korea.

